# Cellular cruise control: energy expenditure as a regulator of collective migration in epithelia

**DOI:** 10.1101/2024.05.21.595054

**Authors:** Isaac B. Breinyn, Simon F. Martina-Perez, Ruth E. Baker, Daniel J. Cohen

## Abstract

Epithelial migration is implicit in processes ranging from gastrula development to the healing of skin, and involves the coordinated motion, force production, and resulting energy expenditure of thousands of constitutive cells. However, the spatiotemporal patterning and regulation of energy expenditure during epithelial migration remains poorly understood. Here, we propose a continuum mechanics framework and use it to explore how energy expenditure regulates epithelial migration. We use canonical mechanical metrics such as force, work and power to define what it means for a tissue to migrate ‘efficiently’ and show that freely expanding epithelia actively regulate themselves to operate within a maximally efficient regime. We then leverage electrotaxis (directed motion in response to an externally applied electric field) as a tool to study non-homeostatic migration using this new framework. We show that regulation of migration is robust to external cues and acts to to attenuate a tissues response to stimuli.

## Introduction

Szent-Gyorgyi’s 1937 statement that “without energy…the cellular fabric would collapse” emphasizes both our long history of trying to understand bioenergetics and the fundamental importance of energy production and consumption for keeping living systems running. The regulation of energy production and consumption is vital to all living systems, and this importance motivates a key question in multicellular systems: when and where do cells in a group use energy, and to what extent might they collectively coordinate energy usage? Myriad cellular processes spanning embryonic development, wound healing, and even cancer invasion, involve large groups of cells moving, and movement requires energy. Motivated by the foundational importance of such collective cell migration, and leveraging the intrinsic connection between cell motility and power consumption, we investigate the relationship between collective migration in large cellular groups and the patterns of power generation needed to drive this motion.

We focus on the collective motion of epithelia–sheets of highly collective cells with strong cell-cell adhesion that play key roles in multicellular development and function [1]. Collective cell migration is central to epithelial function as it allows epithelia to move, heal, and maintain barrier function. Emergent migration patterns within epithelia include domains of highly correlated collective motion and intricate leader-follower dynamics [2–4]. In all cases, cell-cell mechanics play a critical role [5–9], with several studies demonstrating that tightly coordinated patterns of force production in a wide range of tissues are both the cause and consequence of morphogenetic and migratory events [2; 10–15].

As all cell migration ultimately requires cells to exert energetically costly active forces, it seems likely that the coordinated patterns of collective migration might reflect underlying, emergent patterns of coordinated power generation and energy consumption, and raises classic questions of collective behavior where some cells may contribute more, energetically speaking, to the collective motion than others. Answering these questions requires understanding the organization of forces and their resulting mechanics in collective epithelial migration, and has been previously studied in three ways. First, significant work has been done to experimentally relate spatiotemporal patterns of cell velocities across tissues to the underlying traction forces and cell-cell tensile forces, and this work demonstrates that there is substantial heterogeneity [6–8; 16–18]. Second, computational metrics such as entropy, enthalpy, and vorticity have been used to describe abstract concepts of energy in migrating monolayers through the lens of statistical mechanics and fluid dynamics [14; 16; 19]. Finally, molecular mechanism studies have drawn connections between cell metabolism and migration via experimental measurements of ATP/ADP and NADH/NAD^+^ ratios [20; 21], mitochondrial proximity to cellular motor proteins [22], and glucose uptake [20], which have revealed both that collective migration is *energetically costly* and that cell metabolism is non-uniform across tissues [20; 21; 23]. While each of these approaches reveals important and unique aspects of bioenergetics, none of them provide a holistic, *physical* view of power generation as a direct cost of collective migration–*e.g.*, relating mechanical power (Watts) to cellular deformation (Joules) and cell migration velocity (*µ*m/hr).

We hypothesised that the energy expended during collective migration is related to, and regulated by, the propagation of mechanical stresses. To explore this hypothesis, we collected new experimental data of collective epithelial cell migration and then used a continuum modeling approach to compute *mechanical work*. Mechanical work–the energy costs of the forces produced by a migrating monolayer–is a powerful concept for collective migration because it allows the use of continuum models to connect experimental patterns of collective cell migration and deformation to the energy that was required to produce those patterns via active force generation by cells within the tissue. By relating the average rate of mechanical work in a monolayer to its collective migration, we discovered a possible regulation mechanism that reflects an ‘optimal rate of energy expenditure’, above which ‘high spending’ regions of the tissue are decelerated, and ‘low spending’ regions are accelerated. We show that this regulation occurs persistently, dynamically, and precisely in freely expanding tissues and those subject to external stimuli that drive faster migration.

We mapped the energetic performance landscape of collective migration by forcing tissues to migrate far from equilibrium. We achieved this by using direct current electric fields to induce collective electrotaxis–the directed migration of cells in response to physiological ionic currents–the only known cue that can precisely and rapidly trigger collective epithelial migration [24–27]. By using both strong and weak electric fields, we were able to comprehensively characterize the spatiotemporal dynamics of energy expenditure during collective epithelial migration under external control. Our results support that this regulatory mechanism is robust across different forms of collective migration.

## Results

### Quantifying mechanical energy expenditure during migration

We first collected experimental data on a standard epithelial line–the MDCK renal epithelial model. These cells are well-known for their consistent and complex collective dynamics resulting from strong cell-cell adhesions [28], and they rank among the most deeply studied collective tissues as a result [29]. We engineered arrays of 5×5mm^2^ MDCK-II monolayers (approx. 25,000 cells) and allowed them to freely migrate and grow using our established stencil-patterning method (Methods and Supplementary Movie 1) [14; 15; 30; 31]. We then performed time-lapse microscopy using phase contrast imaging (Figure 1a and Methods). We used particle image velocimetry (PIV) to analyze these movies and extract velocity vector fields describing the migration patterns within the tissues. These patterns, shown in Figure 1b, can be effectively visualized using line-integral convolutions of the vector fields where color represents speed and texture represents flow dynamics (Figure 1a, Supplementary Movie 2). The flow dynamics shown in (Figure 1a) show how migration in the bulk (center) of an epithe-lium exhibits random fluctuations while cells at the edges show highly directed motion into the surrounding free space [14; 32]. The random fluctuations in the bulk are consis-tent with previous studies, which report that collective migration patterns in epithelial tissues are spatiotemporally heterogeneous, with domains of coordinated cell motility on the order of 150-300 *µ*m (∼7-15 cells wide) [7; 14; 16; 30; 33–35]. To understand how these patterns of motion incur energy expenditure, we used continuum mechan-ics modelling to relate observed velocities with cell tractions and energy expenditure, defined as the mechanical work done by the tissue during collective migration.

**Figure 1:**
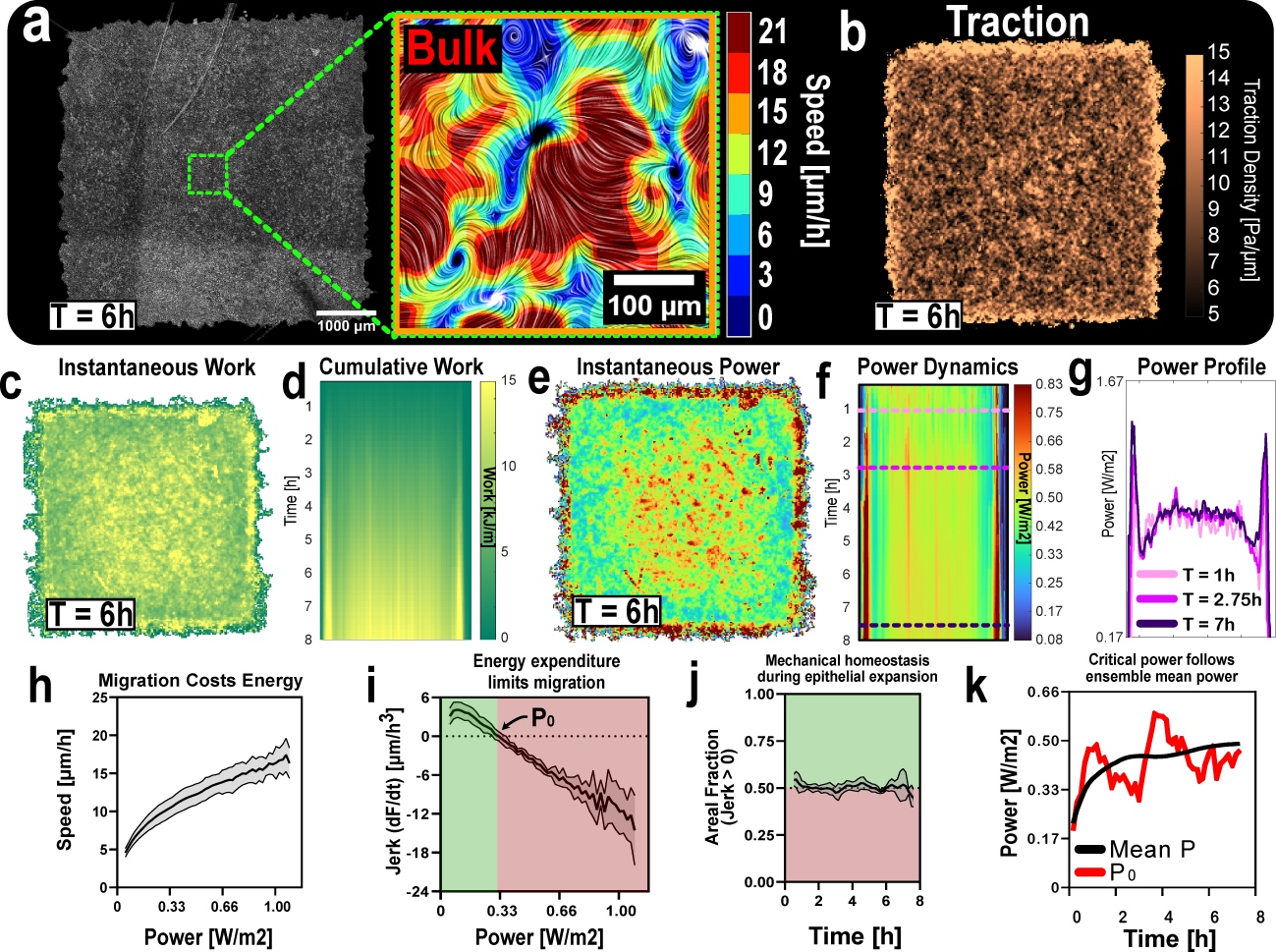
Relating mechanical behaviors to predicted energetic costs in growing epithelial monolayers. **(a)** A phase image of a representative 5×5mm^2^ MDCK-II monolayer 6h post stencil pull. (Green) A 300×300*µ*m^2^ region at the center of the tissue with motion texture produced by LIC and color representing speed [*µ*m·h*^−^*^1^]. **(b)** Heatmap of the computed magnitude of the tractions in all averaged tissues at 6h post stencil pull. **(c)** Heatmap of cumulative work in all averaged tissues.**(d)** Kymograph of cumulative work in all averaged tissues. **(e)** Heatmap of computed power in all averaged tissues. **(f)** Kymograph of power in all tissues averaged. **(g)** Cross-section of power kymograph at different time points (see corresponding lines in **(f)**). **(h)** Velocity increases monotonically with energy expenditure, *i.e.* power. **(i)** Change in force, *i.e.* jerk, changes sign at a critical power, *p*_0_. **(j)** The mean areal fraction of positive jerk, averaged over all tissues. **(k)** Temporal dynamics of ensemble-averaged power and the critical power, *p*_0_.

As the mechanical work relies on force and displacement, we first analyzed patterns of force. Prior studies have used traction-force microscopy [5; 6; 8], and the resulting cell-substrate force measurements inform models predicting cell-cell tensions [7; 36–38]. These tractions, described by a vector field, **T**, propagate through tightly coupled intercellular adhesions [39–41], creating local stresses that can be represented using the Cauchy stress tensor, ***σ***. The relationship between these quantities depends on material properties and has been well characterized by continuum mechanics models in the context of epithelia [42–46], meaning that stresses and tractions within an epithelial monolayer can be well predicted based on the migration velocity field. Mechanically, an epithelial monolayer behaves like an active and incompressible viscous fluid [46]. Given this constitutive law, the Cauchy stress tensor, ***σ***, satisfies ***σ*** = 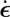, where ***ɛ*** is the strain tensor, ***ɛ*** = (∇**u** +∇*^T^* **u**)*/*2, and **u** is the displacement field so that **v** = 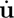. A well-established continuum model for cell-substrate frictions [5; 46] relates velocities and tractions using conservation of linear momentum,

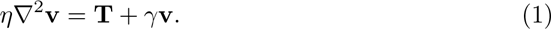

The left-hand side of Equation (1) represents a viscous force per unit area that depends on the viscosity, *η >* 0, and opposes spatial variations in velocity. The right-hand side encapsulates active cell tractions, **T**, and a friction force per unit area that is linear in the cell velocity and depends on the friction coefficient, *γ >* 0 [36; 37; 44; 47].

Combining the continuum framework in Equation (1) (Supplementary Information) with PIV migration data allows us to predict continuum tractions in freely expanding epithelial monolayers. Figure 1b shows a heatmap of the magnitude of the resulting traction field at *t* = 1 hour. In the bulk of the tissue, the magnitude of the computed tractions oscillates both in space and in time (Supplementary Movie 3), consistent with previous work showing that forces propagate through the tissue as waves [48; 49]. The edges, in contrast, exhibit an uninterrupted front of almost-constant traction force magnitude (Figure 1b); consistent with experimental data [17; 41; 48].

Active materials, such as migrating epithelia, incur energetic costs as they exert forces and operate out of thermodynamic equilibrium. A continuum framework allows evaluation of the energetic costs incurred as a result of the mechanical work done by the tissue during migration. The mechanical work per unit area, *W*, depends on the traction force, **T**, and velocity, **v**, as *∂W* (*x, t*)

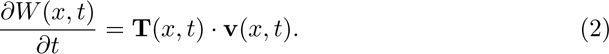

Collective migration is characterized by a flow that creates large deformations in the tissue, *i.e.* the cells that occupy one spatial location at some time point will occupy another spatial location at a future time point. To account for this advection, we use the material time derivative, D*/*D*t*, to calculate the work done by each specific tissue element (Supplementary Information). A spatial distribution of work per unit area at *t* = 8 hours is shown in Figure 1c, with regions of high work per unit area localized to the bulk and edges of the tissue where viscous dissipation and friction forces dominate, respectively (Supplementary Movie 4). We represent the dynamics of the work per unit area (J/m^2^) using a kymograph (Figure 1d, Methods). This quantity does not allow us to compare energetic outputs between tissue regions at different time points directly, so we also calculate mechanical power (work per unit time; W/m^2^), averaged over the lifetime of each tissue region (see Supplementary Information) as a measure of the instantaneous energy expenditure. The spatial distribution of mechanical power is visualized using a heatmap at *t* = 6 hours (Figure 1e). This heatmap shows that the average mechanical power is highest and constant at the tissue edges, lowest in the region between the edges and the bulk, and then increases towards the bulk of the tissue (Supplementary Movie 5). Strikingly this spatial distribution of power is conserved throughout the 8-hour experiment (Figure 1f,g), consistent with experimental data showing that edge expansion is constant in freely expanding MDCK epithelia [14].

Our key finding here is that the energetic costs of collective migration are spatially patterned. That high power regions can emerge from either ‘productive’ migration (*e.g.* outwards expansion and growth) or ‘less productive’ migration (*e.g.* large fluctuations in the bulk without net translation) suggests that the concept of ‘efficiency’ may help with interpreting power consumption data, as we discuss later.

### Epithelial migration is regulated by the rate of energy expenditure

This new analytical framework enabled us to investigate mechanical energy costs as a function of migration patterns. Firstly, we found a monotonic relationship between energy expenditure and migration speed (Figure 1h). The large correlation length in epithelia produces domains of coordinated cell motility that can accelerate relative to neighboring zones [49], dividing the tissue into heterogeneous regions of high and low migration speed. *A priori*, this heterogeneity seems inefficient for net growth relative to a uniform distribution of energy costs, raising the question of whether there is a mechanism that attenuates force production in ‘high-spending’ regions. To investigate this, we sought to relate the average mechanical power to the rate of change of traction forces (d**T**/dt; *i.e.*, jerk). We plotted jerk as a function of mechanical power averaged across tissues (Supplementary Information). Strikingly, we found that the jerk is directly related to average power: future force production increases (decreases) when average power is below (above) a *critical power*, *p*_0_ (Figure 1i), suggesting that the collective is somehow regulating its power consumption around a set point.

We expected that this regulation mechanism would play a role in the proportion of the tissue that could increase speed at any given time. Namely, by symmetry, we expect that at steady-state, approximately half of the tissue would generate more tractions than the critical power dictates, therefore accelerating with respect to the tissue average, and *vice versa*. To test this, we calculated the fraction of the tissue area which was accelerating as a function of time. The fraction of tissue area that is accelerating remains at 0.5, as shown in (Figure 1j) throughout the experiment. These results suggest that the critical power, *p*_0_, is not static. For each time point, we computed the spatial mean of the mechanical power, ⟨*P* ⟩ and the critical power, *p*_0_. A comparison of the two (Figure 1k) shows that the critical power, *p*_0_, oscillates around the average power, ⟨*P* ⟩, with a period of approximately 1-2 hours. This suggests that *p*_0_ is continuously adjusted to restore the fraction of the tissue with positive jerk to its homeostatic value of 0.5.

### Driven collective migration is constrained by energetic costs

Having established a framework that relates tractions and average mechanical power in freely expanding epithelial monolayers, we next explored how the joint dynamics of force and power change when epithelia are driven by an external stimulus. Here, we chose electrotaxis due to its near universality [24; 50–52] and ability to be precisely triggered [27; 53]. Motivated by the range of physiologically relevant field strengths recorded *in vivo* [54], we used two different voltage traces: a 3 V/cm pulse, and a 1.5 V/cm pulse (voltage traces shown in (Figure 2b)).

**Figure 2:**
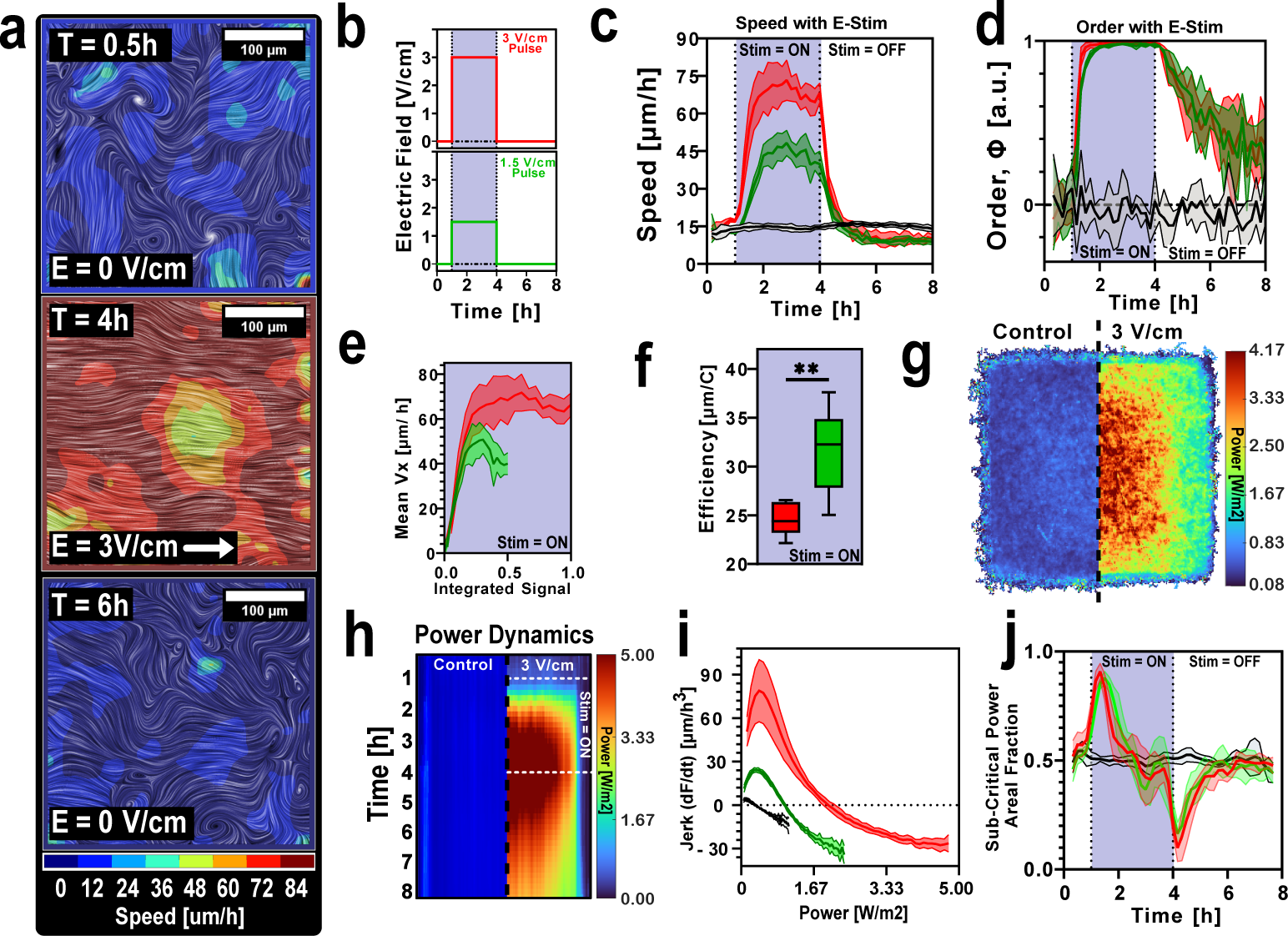
Energetics of externally driven epithelial migration. **(a)** A 300×300*µ*m^2^ region at the center of an electrotaxing tissue with motion texture produced by LIC and color representing speed [*µ*m h*^−^*^1^]. **(b)** The two different stimulation traces used for the remainder of this study. **(c)** Speed dynamics in electrotaxing tissues. **(d)** Order dynamics in electrotaxing tissues. **(e)** Speed as a function of integrated signal (integrated area of signals shown in **(b)**). **(f)** The efficiency of different stimulation traces, expressed as mean cell displacement per unit of injected charge. **(g)** Heatmap of power averaged over all control tissues (left) and all tissues stimulated with the 3 V/cm pulse trace (right). **(h)** Power kymograph averaged over all control tissues (left) and all tissues stimulated with the 3 V/cm pulse trace (right). **(i)** Rate of traction (jerk) vs. power for the different stimulation traces. **(j)** Areal fraction of positive jerk averaged over all tissues for different stimulation traces.

Collective electrotaxis is typically quantified by tracking changes to the mean speed and directional order [25; 27; 30; 53]. The directional order of the tissue, Φ, is defined as

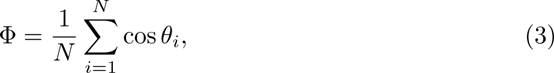

where *θ_i_*is the angle between the velocity vector at location *i* and the electric field, and *N* is the number of velocity vectors. Given the non-uniform response of MDCK tissues to electrotaxis, we focus our remaining analyses on the 2×2mm^2^ bulk region of the tissues. Induced speed in the tissue bulk is shown in Figure 2c. With a 3 V/cm pulse, tissues exhibit a dramatic acceleration in the first hour of stimulation before plateauing and even slowing down *during* stimulation, as shown in previous publications [26]. With a 1.5 V/cm pulse, tissues exhibit a lower peak speed, and a nearly identical slowdown in the latter half of stimulation. Despite these differences in induced speed, the dynamics of the directional order parameter are similar, with both stimulation conditions resulting in strongly directed migration (Figure 2d).

Electrical stimulation enables very clear mapping of the input stimulus (total charge injected) to the output response (Figure 2e–the total stimulation signal is normalized to 3 V/cm, the peak stimulus used). The dynamics here show only marginal gains in speed after increasing the field strength, which provokes a consideration of the efficiency of the electrotactic response as a function of the total stimulus delivered. Defining migratory efficiency during electrotaxis as the displacement of the constituent cells within the tissue per unit of injected charge revealed that the 3 V/cm pulse stimulation protocol was less efficient (Figure 2f), despite producing higher peak speeds.

Given the dramatic impact of electric fields on collective migration, which we expect to be associated with increased force production [55] and energy expenditure, we sought to use our new analytical framework to understand how the relationship between average mechanical power, tractions and velocities is affected during electro-taxis. We first quantified the average power (Figure 2g), comparing control tissues to electrotaxing tissues after *t* = 2 hours of electrotaxis for a 3V/cm pulse (all conditions shown in Supplementary Movie 8). The power heatmap shows that the bulk of the tissue is most active during electrotaxis, which is consistent with prior studies showing the tissue bulk develops highest electrotactic speeds [30; 56]. This result is further corroborated by the kymograph in Figure 2h, which shows an increase in power consumption shortly after stimulation begins. The high-power zone in the tissue bulk shown in Figure 2h exhibits an average mechanical power far greater than in the freely expanding case (Figure 1). To investigate whether mechanical power is regulated during electrotaxis, we calculated the jerk as a function of power (Figure 2i). Once more we see the existence of a critical power beyond which the jerk is negative in both stimulation cases (Figure 2i), with *p*_0_ scaling with the electric field strength. This demonstrates that an external stimulus, such as an electric field, can shift an epithelium’s regulatory behavior.

We then computed the fraction of the tissue that was accelerating over the entire course of the experiment. Both stimulation protocols induce an initial rapid increase in the areal fraction of positive jerk, and tissues return to an areal fraction of 0.5 approximately halfway through the stimulation period (Figure 2j). When stimulation ceases there is a rapid drop in the areal fraction, before a return to baseline. While the tissues rapidly return to an areal fraction of 0.5 during electrotaxis, their mean areal fraction during the experiment is higher than that of an epithelium not undergoing electrotaxis (Supplementary Figure 1).

### Joint dynamics of migration and energy consumption

Having understood that energy consumption actively regulates force production, we now turn our attention to the resulting joint dynamics of energy consumption and velocity. We do this by computing empirical two-dimensional phase portraits of velocity and power (Methods), where streamlines represent trajectories within that space (Figure 3). Since the electric field is in the *x*-direction, we specifically analyze the *x*-component of the velocity. These phase portraits graphically represent the evolution of velocity and power and allow us to interrogate and interpret any feedback between these quantities. The phase portraits for all control tissues (Figure 3a) show that trajectories diverge from the *v* = 0 line towards higher speed and power. This phase portrait additionally reveals a global trend in which trajectories are restored to-wards lower speeds and subsequently lower energy expenditures. Inducing electrotaxis resulted in a striking difference in the phase portrait relative to the unstimulated con-trol tissues (3 V/cm in Figure 3b, 1.5 V/cm in Supplementary Figure 2). Firstly, the phase portraits generated during electrotaxis span a substantially larger domain than the phase portraits for a control tissue. Secondly, despite the electric field providing a constant cue for directed migration, the phase portrait reveals a global decrease in velocities past a certain power.

**Figure 3:**
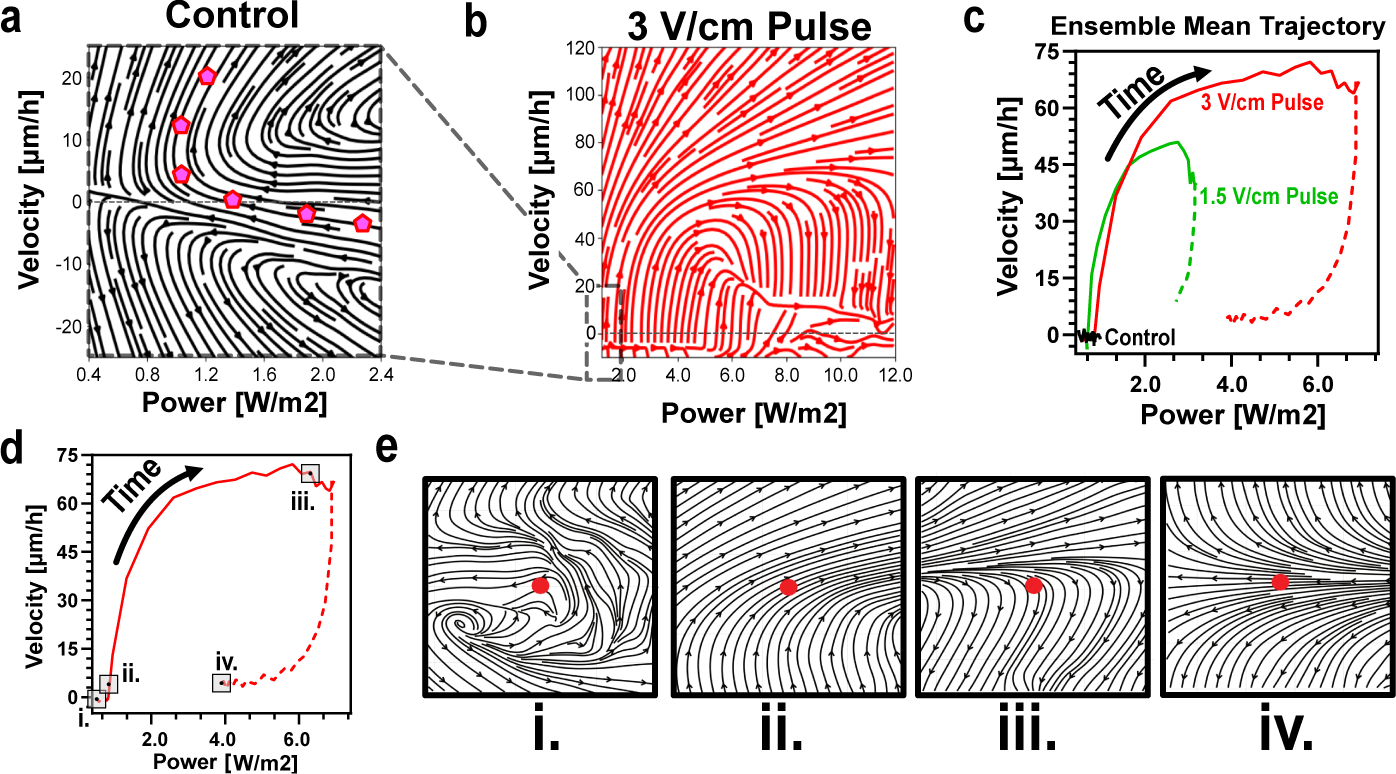
Dynamic behavior of velocity and power within a collectively moving epithelium. **(a)** *v*-*p* space portrait averaged over all control tissues with a representative trajectory shown using pink pentagons. **(b)** *v*-*p* space portrait averaged over all tissues stimulated with a 3V/cm pulse. **(c)** Trajectory of the ensemble mean over time across all experimental conditions. **(d)** Timepoints at which we calculate higher resolution *v*-*p* space portraits. **(e)** Higher resolution *v*-*p* space portraits.

To visualize ensemble behavior, we plotted average trajectories in the *v*-*p* space, which show how energy expenditure rapidly increases during stimulation (solid lines) and rapidly decreases when stimulation is ceased (dashed lines, Figure 3c). To understand the effect of the regulation mechanism on the ensemble velocity, we choose four representative time points at which to probe the structure of the phase portrait at a higher resolution: the beginning of the experiment, the first hour of stimulation, the final hour of stimulation, and the end of the experiment (Figure 3d). When observing the phase portrait immediately surrounding the collective mean at the start of the experiment, we see highly random movement in phase space (Figure 3e(i)). This is consistent with how the ensemble of cells in the center of the tissue responds in the period immediately after stencil removal where the tissue begins outwards motion. At the start of stimulation (Figure 3e(ii)), the collective mean power exists within a uniform region of the phase space in which all states are driven towards a higher power and higher velocity. However, in the final hour of stimulation (Figure 3e(iii)), a bifurcation occurs: some cells continue to accelerate while others begin to slow down, despite the stimulation. We will explore this distinction, and the possibility of different energetically active populations, in the subsequent section. Finally, several hours after stimulation has stopped (Figure 3e(iv)) the phase portrait appears to resemble that in the control tissue.

### Cell-level power regulation and sub-populations

So far, the relationship between power and velocity has been explored at the continuum scale; to get a more granular understanding of these dynamics, we next explore dynamics at the level of individual cells by measuring every cell trajectory (Methods) and computing the average mechanical power per cell over time (Supplementary Information). This allowed us to divide the population of cells into two groups: *weak* cells whose power is *less* than the mean power of the monolayer at *t* = 1 hour of electro-taxis, and *strong* cells whose power is *greater* than that power. Figure 4a shows that the *strong* cells exhibit higher velocities than the *weak* cells in both stimulation cases, with the *strong* cells in the 3 V/cm pulse having the highest velocity. However, over time *strong* cells exhibit a dramatic decrease in speed, with their velocities converging with those of *weak* cells. In contrast, weak cells exhibit a gradual rise to their steady-state speed. In the case of the 1.5 V/cm pulse, the separation of speeds between *strong* and *weak* cells is less pronounced.

**Figure 4:**
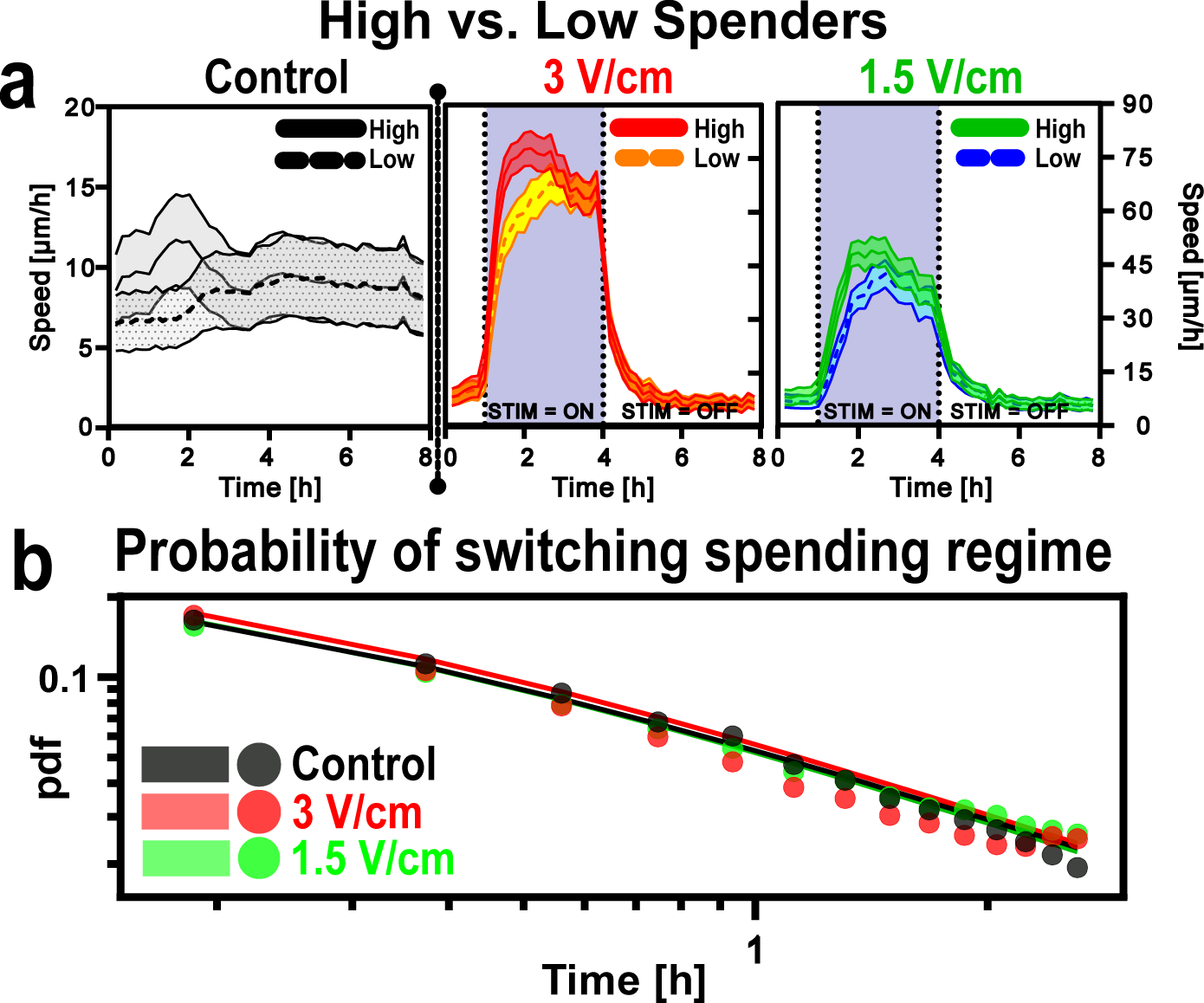
High and low power cells during collective migration. **(a)** Speed dynamics of ‘strong’ (solid lines) and ‘weak’ (dashed lines) cells in tissues exposed to no field (left), a 3 V/cm pulse (middle), and 1.5 V/cm pulse (right). **(b)** Probability density functions of switching spending regime (high to low, or vice versa) for all experiments. Points represent data, solid lines represent power law fits. Please note important changes in axis limits in panel **(a)**.

To explore whether cells are intrinsically *strong* or *weak*, or if this identity is dynamic, we computed the empirical probability distribution of the time that cells remain in a high-spending or a low-spending state, *i.e.* a *residence time* (Supplementary Information). Strikingly, all distributions, regardless of electric field strength, follow the same pattern (Figure 4e), suggesting that the likelihood of a cell switching its spending state is independent of the strength of stimulation. To better understand the empirical distribution of these residence times, we fitted a power law and an exponential law to the residence time data (Supplementary Figure 3), which revealed that the power law, and not the exponential law, fits the data well. This suggests that switching between a high and a low spending state is not merely a stochastic process, but has an inherent memory.

### Regulation is dynamic in time and adapts to external stimuli

Common stimuli such as chemical gradients elicit a time-dependent response, with cells habituating to the stimulus and weakening their sensitivity over time [57; 58]. During electrotaxis, a similar phenomenon occurs (Figure 1c), with tissues slowing down during persistent stimulation. We wondered whether a change in the sensitivity to the electric field would also be apparent in the *v*-*p* phase portraits (Figure 3). To probe this, we computed phase portraits for different one-hour intervals, averaged over all tissues stimulated with a 3 V/cm pulse (Figure 5a) and all tissues in control conditions (Figure 5b, 1.5 V/cm in Supplementary Figure 4).

**Figure 5:**
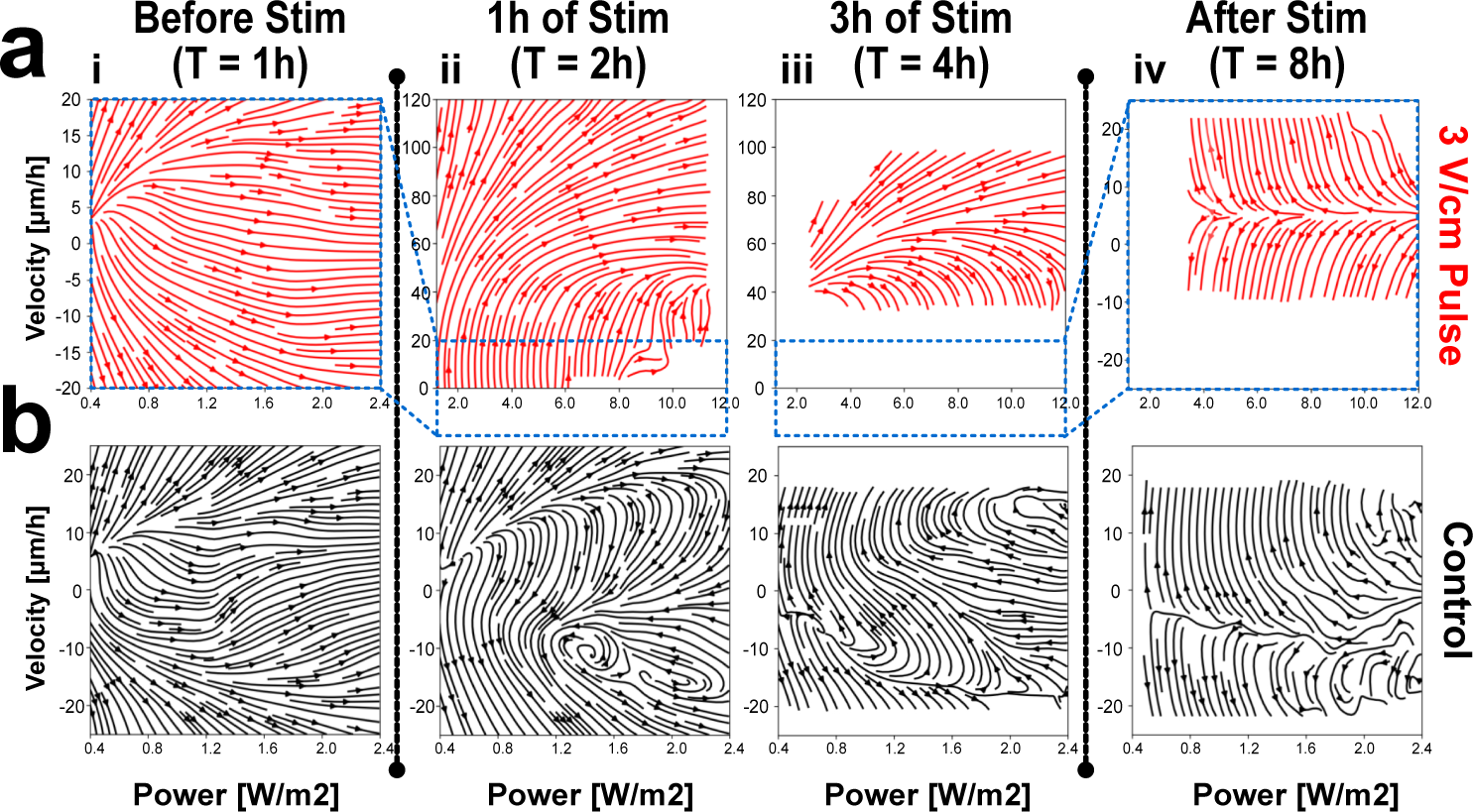
Dynamics of regulation during an external stimulus. **(a)** Phase portraits for one-hour intervals averaged over all 3 V/cm stimulated tissue. (i) first hour of the experiment (no electric field); (ii) first hour of electrotaxis; (iii) third (last) hour of electrotaxis; (iv) two hours post-stimulation. **(b)** Phase portraits for control tissues (no electric field) over the same one-hour intervals.

During the hour before stimulation, the phase portraits are indistinguishable across the different experiments (Figure 5a(i),b(i)). During electrotaxis, the joint dynamics of velocity and average mechanical power not only depart dramatically from those in control conditions but are also far from time-autonomous. While the phase portrait in the first hour of electrotaxis (Figure 5a(ii)) is similar to the average phase portrait for the full three-hour electric field stimulation window (Figure 3b), the *v*-*p* dynamics in the last hour of electrotaxis (Figure 5a(iii)) are characterized by a region in the phase portrait where the entire tissue is slowing down, regardless of average mechanical power. No such effect is observed in the phase portraits in the control experiment during the same time intervals (Figure 5a(ii)-(iii)). Finally, the phase portrait in the fourth hour after the electric field is turned off (Figure 5a(iv)) is again indistinguishable from that of control conditions (Figure 5b), showing a complete return to control conditions. Together, these data suggest that the regulation of active force expenditure varies over time, suggesting potential adaptation in the transduction process (see Discussion).

## Discussion

Here, we proposed a method to quantify the *physical* energy expended during collective cell migration in terms of the mechanical properties of the monolayer. We utilized a continuum model for stress in an epithelial monolayer to calculate mechanical metrics, including force, work and power from empirically measured velocity fields. We found that cells appear to regulate how strongly they contract against the substrate based on their energy expenditure relative to some *critical power*. This regulation remains evident during electrotaxis, with the *critical power* increasing as a function of electric field strength.

A key practical finding of our data is that while electrotaxis significantly boosts collective cell migration speed (∼2-6×) it carries an even larger increase (∼10×) in the energetic costs. This increased energetic cost raises two key questions. First, can these costs be linked to true metabolic costs, which may be rate limiting? A limitation of our study is that we cannot directly measure these costs, but future studies may focus on directly measuring metabolic activity to test these hypotheses [20; 21; 59]. Second, can consideration of energetic costs and their regulation inform intelligent control of electrotaxis for practical use cases *e.g.* wound healing? Notably, we found that weaker electric fields induced more efficient, albeit slower, migration (in terms of displacement per unit of injected charge and energy expenditure). Ultimately, understanding the relationship between the applied field, energy, and induced speed can help us to design more effective stimulation strategies.

Finally, the finding that regulation of active force expenditure is not time-autonomous but changes as the monolayer is stimulated with the electric field could imply that cells within the monolayer adapt to the field, prompting questions about the nature of collective signal transduction within the monolayer. At the same time, the reduction in induced speed within the monolayer could be due to mechanical changes within the tissue, which deforms and possibly sees structural changes during migration. Ultimately, the finding that the regulation mechanism between energy and velocity within the monolayer evolves as the monolayer is exposed to the electric field provides a starting point for future work focused on a combination of signal transduction and mechanics.

## Materials and Methods

### Cell culture

Wild-type Madin-Darby Canine Kidney II cells (MDCK-II, courtesy of the Nelson Lab, Stanford) were cultured in media consisting of low-glucose (1 g/L) DMEM with phenol red (Gibco; powder stock), 1 g/L sodium bicarbonate, 1% streptomycin/penicillin (Gibco), and 10% fetal bovine serum (Atlanta Biological). Cells were maintained at 37°C and 5% CO2 in humidified air. All experiments were conducted between P12 and P18 and cells were mycoplasma tested twice annually.

### Microscopy

For all experiments, time-lapsed montage images were captured every 10 minutes for 8 hours with an automated inverted microscope (Zeiss Axio Observer Z1) equipped with a 5x/0.16 phase contrast Zeiss objective, a Photometrics Iris 9 (Photometrics, Inc.) sCMOS camera, and an XY motorized stage, all controlled using Slidebook (3i Intelligent Imaging Innovations). Additionally, the stage-top was enclosed in a cage incubator kept at 37 °C. Captured images were stitched using (Fiji Is Just) ImageJ’s ‘Grid/Collection stitching’ plug-in (three 3×3 montages w/ 15% spatial overlap) [60], template matched using the ‘Template Matching’ plug-in, masked, and aligned before further analyses.

### Electrotaxis assays

The bioreactor used for electrotaxis was modified from our SCHEEPDOG platform for uniaxial stimulation of large-scale tissues, and is described in complete detail in previous works from the lab [30; 31; 53; 56; 61; 62]. Briefly, a laser-cut and glued acrylic body houses an electrode (Ag|AgCl) -electrolyte (PBS) -salt bridge (4% w/v agarose/PBS) assembly and is clamped to a 10 cm tissue culture dish to create a microfluidic region supported by a 250 *µ*m thick stencil of PDMS. Electrodes protrude from the top of the device, allowing for direct integration with external controllers and the body contains both an inlet and outlet for fresh media perfusion orthogonal to the direction of electrical stimulation; this aids in temperature regulation (see [31] for full characterization) and perfusion of electrochemical byproducts such as reactive oxygen species. Titanium probes are inserted into the agarose salt bridges to monitor the voltage and current across the microfluidic channel, measured by a USB oscilloscope (Analog Discovery 2, Digilent Inc.) and proportionally controlled by a Keithley 2450 SourceMeter (Tektronix) and custom MATLAB script. For a complete summary of the approach and guidelines, CAD files of the full device, and/or custom closed-loop control scripts, see previous works [30; 31; 53; 56; 61; 62] and our repositories (github.com/cohenlabprinceton).

### Tissue culture

Per experiment, three 5×5 mm^2^ square tissues were seeded onto 10 cm tissue-culture plastic dishes (CELLTREAT) using stencils of 250 *µ*m-thick silicone elastomer (Bisco HT6240) cut with a hobbyist razor-writer (Silhouette Cricut). Suspensions of MDCK cells were seeded into the stencil patterns at a volume of 10 *µ*L and density of 2.5 (±0.15) x 10^6^ cells/mL. Cells were given 1 hour to adhere before the dish was flooded with media and incubated for 18 hours before stencil pull and device integration, as described below.

### Experiment design

In all experiments (with and without electrotaxis), tissue culture stencils were pulled approximately 1 hour before imaging began. Immediately post-stencil pull, dishes were refilled with fresh culture media followed by direct integration with the bioreactor. The assembled bioreactor is then placed into the microscope (see Microscopy), situated within a custom-built cage incubator maintained at 37 °C, and perfused with fresh media continuously bubbled with 5% CO_2_ using a peristaltic pump (Instech Laboratories) at a rate of 2.5 mL/hour.

When integrating electrical stimulation, experiments begin with a 1-hour ‘control’ period before stimulation is turned on. Relevant stimulation is then delivered for 3h (see Bioreactor) before a 4-hour ‘relaxation’ period during which there is no stimulation but imaging persists.

### Fluorescence microscopy reconstruction

All nuclear data used for computational analyses were calculated from transmitted light phase contrast microscopy using our in-house Fluorescence Reconstruction Microscopy tool (convolution neural network trained to produce images of various biomarkers from phase contrast images) [33].

### Kymographs

To calculate work and power kymographs, MxNxT heatmap stacks are first calculated for the corresponding metric. Then, a 2 mm x N rectangle is cropped from the center of the tissue and averaged along the first axis, resulting in a one-dimensional vector of size 1xN. This is then repeated for all timepoints, T, with the resultant vectors populating a TxN kymograph. After population, a moving mean of window size 3 (30 minutes) is applied along the vertical (temporal) axis.

### Particle image velocimetry and particle tracking

After initial image preparation (see Microscopy), velocity vector fields were calculated using MATLAB’s PIVlab [63]. A 2-pass fast Fourier transform was performed with interrogation window sizes of 128 x 128 pixels and 64 x 64 pixels with 50% overlap. Vectors outside of five standard deviations were filtered out and replaced with interpolated data. After conversion of phase contrast data to nuclear data (see Fluorescence microscopy reconstruction), we used ImageJ’s TrackMate [64] to track all cells throughout the experimental time course (tracks shorter than two thirds of the entire experiment were filtered out).

### Phase portrait calculations

Using the measured velocity data and the computed time-averaged mechanical power, empirical two-dimensional phase portraits of velocity and power can be computed. Since the electric field is in the *x*-direction, we analyze the *x*-component of the velocity, which we denote by *v*. The time-averaged mechanical power is computed as before. These phase portraits show the *v*-*p* trajectories found in the experimental data set. For the computation of each empirical phase portrait, we consider a dataset that consists of points within a given spatiotemporal domain, *e.g.* all spatial locations between *t* = 1 hour and *t* = 3 hours for a 3 V/cm pulse and we use the same temporal domain for all experimental conditions. We then compute the time derivative of the velocities and power using a first-order finite difference scheme. The time derivatives of velocities and power can be used directly to compute velocities within *v*-*p* space, which are then described by a vector field, **R**, defined as

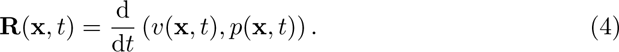

The computed velocities in *v*-*p* space can be used to generate phase portrait plots graphically, using the MATLAB function streamslice [65] or matplotlib function streamplot in Python.

### Averaging phase portraits over biological replicates

Briefly, phase portraits were averaged over biological replicates by first interpolating all first-order time derivatives of velocity and power such that phase portraits of each tissue were sampled on the same grid. The time derivatives of velocity and power were averaged across tissues.

### Energy calculations

All computations are performed using custom MATLAB and Python scripts. Please see the Theory Supplementary Information and Github repositories for code (github. com/cohenlabprinceton).

## Supporting information

Theory SI

SupplementaryMovie1_EpithelialExpansion

SupplementaryMovie2_ControlSpeed

SupplementaryMovie3_ControlTractions

SupplementaryMovie4_ControlWork

SupplementaryMovie5_ControlPower

SupplementaryMovie6_ElectrotaxisFull

SupplementaryMovie7_ElectrotaxisCrop

SupplementaryMovie8_StimPower

## Acknowledgments

The authors would like to thank Yubin Lin for his help running JulieVision. I.B.B. is supported by an NSF GRFP. S.M.-P. is supported by an EPSRC Doctoral Training Award. R.E.B. would like to acknowledge support from the Simons Foundation (grant MPS-SIP-00001828). D.J.C. would like to acknowledge support for this work was provided in part by the National Institute of Health Award R35 GM133574-03 and the National Science Foundation CAREER Award 2046977. R.E.B. and D.J.C. would like to acknowledge support from the Royal Society via the International Exchange Scheme programme.

## Author contributions

I.B.B. performed experiments. I.B.B. and S.M.-P. conceived the project and analyzed data. I.B.B., S.M.-P., R.E.B., and D.J.C. designed the research and wrote the article. R.E.B. and D.J.C. funded and supervised the work. The co-first authors, I.B.B. and S.M.-P., contributed equally and their names appear in alphabetical order. Both co-first authors agree that either may list themselves first on a CV.

## References

[1] Leys, S. P. & Riesgo, A. Epithelia, an evolutionary novelty of metazoans. Journal of Experimental Zoology Part B: Molecular and Developmental Evolution 318, 438–447 (2012).

[2] Friedl, P. & Gilmour, D. Collective cell migration in morphogenesis, regeneration and cancer. Nature reviews Molecular cell biology 10, 445–457 (2009).

[3] Kabla, A. J. Collective cell migration: leadership, invasion and segregation. Journal of The Royal Society Interface 9, 3268–3278 (2012).

[4] Rørth, P. Collective cell migration. Annual review of cell and developmental 25, 407–429 (2009).

[5] Alert, R. & Trepat, X. Physical Models of Collective Cell Migration. The Annual Review of Condensed Matter Physics is Annu. Rev. Condens. Matter Phys. 2020 11, 77–101 (2020). URL 10.1146/annurev-conmatphys-.

[6] Tambe, D. T. et al. Monolayer stress microscopy: Limitations, artifacts, and accuracy of recovered intercellular stresses. PLOS ONE 8, 1–12 (2013). URL 10.1371/journal.pone.0055172.

[7] Blanch-Mercader, C. et al. Effective viscosity and dynamics of spreading epithelia: a solvable model. Soft Matter 13, 1235–1243 (2017). URL 10.1039/C6SM02188C.

[8] Serra-Picamal, X. et al. Mechanical waves during tissue expansion. Nature Physics 8, 628–634 (2012).

[9] Vincent, R. et al. Active Tensile Modulus of an Epithelial Monolayer. Physical Review Letters 115 (2015).

[10] Cheng, Y., Felix, B. & Othmer, H. G. The Roles of Signaling in Cytoskeletal Changes, Random Movement, Direction-Sensing and Polarization of Eukaryotic Cells (2020).

[11] Fouchard, J. et al. Curling of epithelial monolayers reveals coupling between active bending and tissue tension. Proceedings of the National Academy of Sciences 117, 9377–9383 (2020). URL 10.1073/pnas.1917838117.

[12] Barriga, E. H., Franze, K., Charras, G. & Mayor, R. Tissue stiffening coordinates morphogenesis by triggering collective cell migration in vivo. Nature 554, 523–527 (2018).

[13] Von Dassow, M. & Davidson, L. A. Variation and robustness of the mechanics of gastrulation: The role of tissue mechanical properties during morphogenesis (2007).

[14] Heinrich, M. A. et al. Size-dependent patterns of cell proliferation and migration in freely-expanding epithelia. Elife 9, e58945 (2020).

[15] Heinrich, M. A., Alert, R., Wolf, A. E., Košmrlj, A. & Cohen, D. J. Self-assembly of tessellated tissue sheets by expansion and collision. Nature communications 13, 4026 (2022).

[16] Lin, S.-Z. et al. Universal statistical laws for the velocities of collective migrating cells. Advanced Biosystems 4, 2000065 (2020). URL https://onlinelibrary.wiley.com/doi/abs/10.1002/adbi.202000065. . https://onlinelibrary.wiley.com/doi/pdf/10.1002/adbi.202000065.

[17] Saraswathibhatla, A., Galles, E. E. & Notbohm, J. Spatiotemporal force and motion in collective cell migration. Scientific Data 7, 197 (2020).

[18] Bauer, A. et al. pyTFM: A tool for traction force and monolayer stress microscopy. PLoS Computational Biology 17 (2021).

[19] Lin, S.-Z., Zhang, W.-Y., Bi, D., Li, B. & Feng, X.-Q. Energetics of mesoscale cell turbulence in two-dimensional monolayers. Communications Physics 4, 21 (2021). URL 10.1038/s42005-021-00530-6.

[20] DeCamp, S. J. et al. Epithelial layer unjamming shifts energy metabolism toward glycolysis. Scientific Reports 10, 1–15 (2020).

[21] Zanotelli, M. R. et al. Energetic costs regulated by cell mechanics and confinement are predictive of migration path during decision-making. Nature communications 10, 4185 (2019).

[22] Madan, S., Uttekar, B., Chowdhary, S. & Rikhy, R. Mitochondria lead the way: mitochondrial dynamics and function in cellular movements in development and disease. Frontiers in Cell and Developmental Biology 9, 781933 (2022).

[23] Chen, Y. & McDonald, J. A. Collective cell migration relies on ppp1r15-mediated regulation of the endoplasmic reticulum stress response. Current Biology (2024).

[24] Zhao, M. et al. Electrical signals control wound healing through phosphatidylinositol-3-oh kinase-*γ* and pten. Nature 442, 457–460 (2006).

[25] Cohen, D. J., Nelson, W. J. & Maharbiz, M. M. Galvanotactic control of collective cell migration in epithelial monolayers. Nature Materials 13, 409–417 (2014).

[26] Wolf, A. E., Heinrich, M. A., Breinyn, I. B., Zajdel, T. J. & Cohen, D. J. Short-term bioelectric stimulation of collective cell migration in tissues reprograms long-term supracellular dynamics. PNAS Nexus 1 (2022).

[27] Zajdel, T. J., Shim, G., Wang, L., Rossello-Martinez, A. & Cohen, D. J. SCHEEPDOG: Programming Electric Cues to Dynamically Herd Large-Scale Cell Migration. Cell Systems 10, 506–514 (2020).

[28] Ilina, O. & Friedl, P. Mechanisms of collective cell migration at a glance. Journal of cell science 122, 3203–3208 (2009).

[29] Zorn, M. L., Marel, A.-K., Segerer, F. J. & Rädler, J. O. Phenomenological approaches to collective behavior in epithelial cell migration. Biochimica et Biophysica Acta (BBA)-Molecular Cell Research 1853, 3143–3152 (2015).

[30] Wolf, A. E., Heinrich, M. A., Breinyn, I. B., Zajdel, T. J. & Cohen, D. J. Short-term bioelectric stimulation of collective cell migration in tissues reprograms long-term supracellular dynamics. PNAS nexus 1, pgac002 (2022).

[31] Zajdel, T. J., Shim, G., Wang, L., Rossello-Martinez, A. & Cohen, D. J. Scheepdog: programming electric cues to dynamically herd large-scale cell migration. Cell systems 10, 506–514 (2020).

[32] Poujade, M. et al. Collective migration of an epithelial monolayer in response to a model wound. Proceedings of the National Academy of Sciences 104, 15988–15993 (2007).

[33] LaChance, J. & Cohen, D. J. Practical fluorescence reconstruction microscopy for large samples and low-magnification imaging. PLoS computational biology 16, e1008443 (2020).

[34] LaChance, J., Suh, K., Clausen, J. & Cohen, D. J. Learning the rules of collective cell migration using deep attention networks. PLoS Computational Biology 18 (2022).

[35] Vedula, S. R. K. et al. Emerging modes of collective cell migration induced by geometrical constraints. Proceedings of the National Academy of Sciences 109, 12974–12979 (2012).

[36] Alert, R. & Trepat, X. Physical models of collective cell migration. Annual Review of Condensed Matter Physics 11, 77–101 (2020). URL 10.1146/annurev-conmatphys-031218-013516. 10.1146/annurev-conmatphys-031218-013516.

[37] Arciero, J. C., Mi, Q., Branca, M. F., Hackam, D. J. & Swigon, D. Continuum model of collective cell migration in wound healing and colony expansion. Biophysical Journal 100, 535–543 (2011). URL https://www.sciencedirect.com/science/article/pii/S0006349510051921.

[38] Pérez-González, C. et al. Active wetting of epithelial tissues. Nat. Phys. 15, 79–88 (2019).

[39] Cohen, D. J. & Nelson, W. J. Secret handshakes: cell–cell interactions and cellular mimics. Current opinion in cell biology 50, 14–19 (2018).

[40] Gómez-González, M., Latorre, E., Arroyo, M. & Trepat, X. Measuring mechanical stress in living tissues. Nature Reviews Physics 2, 300–317 (2020).

[41] Trepat, X. et al. Physical forces during collective cell migration. Nature physics 5, 426–430 (2009).

[42] Tlili, S. et al. Collective cell migration without proliferation: density determines cell velocity and wave velocity. Royal Society Open Science 5, 172421 (2018). URL https://royalsocietypublishing.org/doi/abs/10.1098/rsos.172421. https://royalsocietypublishing.org/doi/pdf/10.1098/rsos.172421.

[43] Tlili, S. et al. Migrating epithelial monolayer flows like a maxwell viscoelastic liquid. Phys. Rev. Lett. 125, 088102 (2020). URL https://link.aps.org/doi/10.1103/PhysRevLett.125.088102.

[44] Christensen, A. et al. Friction-limited cell motility in confluent monolayer tissue. Physical Biology 15, 066004 (2018). URL 10.1088/1478-3975/aacedc.

[45] Fang, C., Yao, J., Zhang, Y. & Lin, Y. Active chemo-mechanical feedbacks dictate the collective migration of cells on patterned surfaces. Biophysical Journal 121, 1266–1275 (2022). URL https://www.sciencedirect.com/science/article/pii/S0006349522001527.

[46] Vazquez, K., Saraswathibhatla, A. & Notbohm, J. Effect of substrate stiffness on friction in collective cell migration. Scientific Reports 12 (2022). URL 10.1038/s41598-022-06504-0.

[47] Alert, R., Blanch-Mercader, C. & Casademunt, J. Active fingering instability in tissue spreading. Phys. Rev. Lett. 122, 088104 (2019). URL https://link.aps.org/ doi/10.1103/PhysRevLett.122.088104.

[48] Serra-Picamal, X. et al. Mechanical waves during tissue expansion. Nature Physics 8, 628–634 (2012). URL 10.1038/nphys2355.

[49] Peyret, G. et al. Sustained oscillations of epithelial cell sheets. Biophysical Journal 117, 464–478 (2019). URL https://www.sciencedirect.com/science/article/pii/S0006349519305028.

[50] Du Bois-Reymond, E. Untersuchungen über thierische Elektricität, vol. 2 (G. reimer, 1849).

[51] Reid, B. & Zhao, M. The electrical response to injury: molecular mechanisms and wound healing. Advances in wound care 3, 184–201 (2014).

[52] Cortese, B., Palamà, I. E., D’Amone, S. & Gigli, G. Influence of electrotaxis on cell behaviour. Integrative Biology 6, 817–830 (2014).

[53] Zajdel, T. J., Shim, G. & Cohen, D. J. Come together: On-chip bioelectric wound closure. Biosensors and Bioelectronics 192, 113479 (2021).

[54] McCaig, C. D., Rajnicek, A. M., Song, B. & Zhao, M. Controlling cell behavior electrically: current views and future potential. Physiological reviews (2005).

[55] Cho, Y., Son, M., Ko, U. H., Jeong, H. & Shin, J. H. Electric stimulation alters intercellular stress within the epithelial monolayer. The FASEB Journal 32, 869–3 (2018).

[56] Cohen, D. J., James Nelson, W. & Maharbiz, M. M. Galvanotactic control of collective cell migration in epithelial monolayers. Nature materials 13, 409–417 (2014).

[57] Erban, R. & Othmer, H. G. From signal transduction to spatial pattern formation in E. coli: A paradigm for multiscale modeling in biology. Multiscale Modeling and Simulation 3, 362–394 (2005).

[58] Alon, U., Surette, M. G., Barkai, N. & Leibler, S. Robustness in bacterial chemotaxis. Nature 397, 168–171 (1999).

[59] Zanotelli, M. R. et al. Energetic costs regulated by cell mechanics and confinement are predictive of migration path during decision-making. Nature communications 10, 4185 (2019).

[60] Schindelin, J., et al. Fiji: an open-source platform for biological-image analysis. Nature methods 9, 676–682 (2012).

[61] Shim, G., Devenport, D. & Cohen, D. J. Overriding native cell coordination enhances external programming of collective cell migration. Proceedings of the National Academy of Sciences 118, e2101352118 (2021).

[62] Shim, G., Breinyn, I. B., Martínez-Calvo, A., Rao, S. & Cohen, D. J. Bioelectric stimulation controls tissue shape and size. Nature Communications 15, 1–17 (2024).

[63] Thielicke, W. & Stamhuis, E. Pivlab–towards user-friendly, affordable and accurate digital particle image velocimetry in matlab. Journal of open research software 2 (2014).

[64] Tinevez, J.-Y. et al. Trackmate: An open and extensible platform for single-particle tracking. Methods 115, 80–90 (2017). URL https://www.sciencedirect.com/science/article/pii/S1046202316303346. Image Processing for Biologists.

[65] The MathWorks Inc. MATLAB version: 9.13.0 (R2022b) (2022). URL https://www.mathworks.com.

