## Supplementary material for "Cellular cruise control: energy expenditure as a regulator of collective migration in epithelia": Theory SI

(Dated: May 20, 2024)

### CONTINUUM MECHANICS OF FREELY EXPANDING EPITHELIA

In this work, we are interested in quantifying mechanical metrics from experimental data of collective MDCK migration. MDCK epithelial monolayers make up a flat sheet, which means that the ratio of the height, $h$, to the out-of-plane curvature, $R$, is small, *i.e.*,

$$\frac{h}{R} \ll 1. \tag{S1}$$

This scaling justifies the common assumption in the literature that cell forces upon the substrate and within the monolayer are planar and do not have any out of plane contributions [1]. Since cell movements within a monolayer, which we can quantify using PIV (see Figure 1a in the main text), are tightly coupled through intercellular adhesions, active forces within the monolayer create local stresses, which are transmitted throughout the tissue across a typical length scale [2–5]. In a continuum mechanics formulation, forces in a two-dimensional material, like an epithelial monolayer, are described by a force vector field, $\mathbf{F}$, and monolayer stresses are described by the (Cauchy) stress tensor, $\boldsymbol{\sigma}$, which is a tensor field that describes the state of stress of a deformed material. The cell forces within the monolayer are typically interpreted as the active force transmitted by cells onto other cells, as well as on the substrate, to balance the interstitial stresses inside the monolayer, as well to generate their active movement. It is important to note that a continuum force field in an epithelial monolayer describes a variety of forces that cells exert, including active and passive forces exerted on their neighbours, as well as active traction forces.

Forces and stresses are physically related through force balance, which is captured by the law of conservation of linear momentum. In the context of epithelia, inertial effects can be ignored [6], and so conservation of linear momentum reduces to a force balance condition between monolayer stresses and cell (active) tractions, $\mathbf{T}$ [1], which then reads

$$\frac{\partial \boldsymbol{\sigma}_{\boldsymbol{xx}}}{\partial x} + \frac{\partial \boldsymbol{\sigma}_{xy}}{\partial y} = -\mathbf{T}_x^{\star}, \tag{S2}$$

$$\frac{\partial \boldsymbol{\sigma}_{xy}}{\partial x} + \frac{\partial \boldsymbol{\sigma}_{yy}}{\partial y} = -\mathbf{T}_y^{\star}, \tag{S3}$$

where

$$\mathbf{T}_x^{\star} = \mathbf{T}_x/h, \quad \mathbf{T}_y^{\star} = \mathbf{T}_y/h, \tag{S4}$$

and $\mathbf{T}_x, \mathbf{T}_y$ are the $x$ and $y$-components of the tractions, respectively. Equations (S2) and (S3) represent force balance per unit height in the tissue. By additionally assuming that the monolayer is homogeneous and isotropic, Equations (S2) and (S3) allow for the direct computation of monolayer tractions when the state of stress in the tissue is known. Likewise, when the forces are known, for example through traction force microscopy (TFM), the components of the stress tensor, $\sigma$, can be found by solving Equations (S2) and (S3) as PDEs subject to appropriate boundary conditions – for example, no-slip or free boundary conditions [1].

In fact, Equations (S2) and (S3) are the core mathematical foundation of traction force microscopy (TFM) [1]. Indeed, TFM uses a constitutive equation to determine the traction forces from substrate deformations measured by particle image velocimetry to yield a computed traction field, $\mathbf{T}$.

#### Constitutive assumptions

With a known stress field, forces in the tissue can be directly computed through Equations (S2) and (S3). Physically, the state of stress in the tissue can be directly determined by the state of deformation by introducing a *constitutive assumption* that prescribes the relation between stresses and strains. The continuum mechanics of collectively migrating epithelia are characterised by several well-established canonical models that account for a wide range of physical contexts, like varying levels of proliferation, substrate stiffness and tissue geometries [6–10]. The use of such models is supported by a large body of research establishing the validity and applicability of modelling epithelial monolayers as active viscoelastic materials. Viscoelastic models capture key characteristics of the migrating epithelial monolayer, like stress relaxation, which follow from the physical properties of the constituent cells [3, 9].

In general, such models of viscoelasticity provide constitutive relations for the monolayer stresses and activity that are described entirely by the directly observable physical quantities of strain,

$$\boldsymbol{\epsilon} = \frac{1}{2}(\nabla \mathbf{u} + \nabla^T \mathbf{u}), \tag{S5}$$

and strain rate,

$$\dot{\boldsymbol{\epsilon}} = \frac{1}{2}(\nabla \mathbf{v} + \nabla^T \mathbf{v}), \tag{S6}$$

respectively, where $\mathbf{u}$ is the displacement field, and $\mathbf{v} = \dot{\mathbf{u}}$ is the velocity. Recall that in this work, the displacement and velocity fields are found experimentally using PIV from microscopy images. Appropriate constitutive relations may be given by, for example, the Maxwell, Kelvin-Voigt, or simple viscous models. For a review, see [3, 9]. While these models all give rise to different mechanical behaviours in response to external forces and internal activity [3, 5, 9, 11], their applicability as suitable descriptions of monolayer mechanics in different contexts is well established.

In sum, having constitutive equations that link the directly observable velocity field, $\mathbf{v}$, which is measured using PIV, to the stress tensor, $\boldsymbol{\sigma}$, allows for the direct computation of a continuum traction field through Equations (S2) and (S3). In this view, inferring monolayer stresses and forces from velocities reduces to choosing an appropriate constitutive assumption that most faithfully describes the epithelial system of interest. An MDCK epithelial monolayer, with its typical proliferation and strong cell-cell adhesions, can be effectively described as a freely migrating viscous liquid, such that a constitutive assumption of an active viscous material at low Reynolds number is the most appropriate [6]. Such models take into account the stresses within the monolayer and their direct transmission to the substrate by cell-substrate adhesions. By choosing a simple viscous constitutive law the Cauchy stress tensor is readily expressed in terms of the rate of strain tensor,

$$\boldsymbol{\sigma} = \eta \dot{\boldsymbol{\epsilon}}, \tag{S7}$$

where $\eta$ is the effective viscosity of the material. By then making the canonical assumption that tissue flow is incompressible, *i.e.*,

$$\nabla \cdot \mathbf{v} = 0, \tag{S8}$$

and imposing conservation of linear momentum to find the tractions, through Equations (S2) and (S3), the velocity field is described as a generalised Stokes flow given by

$$\eta \nabla^2 \mathbf{v} = \mathbf{T} + \gamma \mathbf{v}. \tag{S9}$$

The left-hand side of Equation (S9) can be interpreted as a viscous force per unit area that opposes spatial variations in the velocity. The right-hand side expresses cell tractions as an external driving force per unit area, $\mathbf{T}$, and also contains a friction force per unit area with the substrate of the form $\gamma \mathbf{v}$, where $\gamma > 0$. This term, which is linear in the cell velocities, is a canonical approximation for the cell-substrate friction and is analogous to the viscous drag experienced by a low Reynolds number fluid flow over a surface [9, 11], which is likewise represented by linear friction.

#### Mechanical parameters

The mechanical properties of epithelial monolayers on soft substrates are well characterized, and theoretical estimates of key mechanical parameters are supplied by the literature. Here, we report the mechanical parameters we have used in our force computations.

| Parameter | Reference value | Source |
|---|---|---|
| Friction coefficient, $\gamma$ | 100 Pas/$\mu$m$^2$ | [2, 12] |
| Monolayer viscosity, $\eta$ | 25 MPa·s | [2, 12] |
| Hydrodynamic length, $\sqrt{\eta/\gamma}$ | 0.5 mm | Computed |

**Table S1**. Mechanical parameters used for the computation of forces and stresses throughout all experiments.

### CONTINUUM COMPUTATIONS OF MECHANICAL WORK

To assess the energetic cost associated with the active cell forces within the monolayer, a simple model for the energy associated with migration in the monolayer is needed. Previous investigations into continuum descriptions of energetics in cellular collectives have used various energy metrics to relate the velocity field to cellular energetics [13, 14]. For example, Lin *et al.* [13, 14] study the distribution of kinetic energy,

$$E = (1/2)|\mathbf{v}|^2, \tag{S10}$$

and enstrophy,

$$\Omega = (1/2)|\boldsymbol{\omega}|^2, \tag{S11}$$

respectively, where $\boldsymbol{\omega}$ is the vorticity tensor. Their investigations find that the spatiotemporal distribution of energy and velocity in the tissue is stochastic and follows a $q$-Gaussian distribution, which is similar to the empirical distribution of monolayer velocities [13]. Although both kinetic energy and enstrophy provide a good mathematical abstraction for the energy stored in a tissue, these two metrics do not reflect the physical work done by the tissue as a consequence of the forces associated with migration. Since Equation (S9) provides a theoretically justified continuum framework to describe forces from velocity data, the question becomes how to use this framework to derive a metric directly informative of the mechanical energy spent during collective migration in the cell monolayer. Perhaps the simplest metric for the mechanical energy associated with force production is that of mechanical work, $W$. Given a traction field, $\mathbf{T}$, and a velocity field, $\mathbf{v}$, the mechanical work per unit area is readily expressed in the Eulerian frame of reference using the standard expression

$$\frac{\partial W(\mathbf{x},t)}{\partial t} = \mathbf{T}(\mathbf{x},t) \cdot \mathbf{v}(\mathbf{x},t). \tag{S12}$$

As we are interested in the regulation of active forces jointly with mechanical work, we would like to track the work done by a particular tissue element, as opposed to that at a particular fixed location in two-dimensional space. These two concepts are different, since monolayer migration is characterised by a flow that creates advection in the tissue, *i.e.*, the cells that occupy one spatial location at some time point will occupy another spatial location at a future time point. Put differently, the fact that the tissue is migrating means that the work, $W$, if interpreted as a scalar quantity associated with a fixed tissue element, is convected along with the monolayer velocity. Therefore, to track the mechanical work done by the monolayer during the course of its migration, it is important to find an expression for the mechanical work done by the part of the tissue occupying any fixed location $x$ at time $t$ during its history. To take into account the convection with the monolayer velocity, and quantify the work that corresponds to a fixed region of tissue that occupies given coordinates in space and time, we use the material time derivative, $\mathrm{D}/\mathrm{D}t$. For a scalar function, $\varphi$, the material derivative is defined as

$$\frac{\mathrm{D}\varphi}{\mathrm{D}t} = \frac{\partial \varphi}{\partial t} + \mathbf{v} \cdot \nabla \varphi. \tag{S13}$$

Combining Equations (S12) and (S13), yields the material derivative of the mechanical work, $W$,

$$\frac{\mathrm{D}W(\mathbf{x},t)}{\mathrm{D}t} = \mathbf{T}(\mathbf{x},t) \cdot \mathbf{v}(\mathbf{x},t) + \mathbf{v} \cdot \nabla W. \tag{S14}$$

#### Computation of power

By integrating in time, Equation (S14) yields the total mechanical energy used by the region of tissue that is at location $x$ at time $t$, *i.e.*,

$$W(\mathbf{x},t) = \int_0^t \frac{\mathrm{D}W(\mathbf{x},s)}{\mathrm{D}s} \mathrm{d}s. \tag{S15}$$

This is a useful metric to understand how much energy has been expended by different regions of tissue at the same time point. In the context of epithelial migration, cells have processes that allow them to absorb and metabolise energy from their environment continuously. Therefore, a metric that relates to the *rate* of energy expenditure in the monolayer can be used more meaningfully to assess energetic cost for the epithelial monolayer during its migration. To achieve this, we consider the quantity of *average mechanical power*. The average power expended by a tissue element allows us to assess the average rate at which energy has been used by a tissue element during epithelial migration, as a function of force production against the substrate and against local deformations.

Defining average mechanical work would be straightforward if one were to ignore tissue expansion, and assume that the monolayer occupies the same spatial extent during the experiment. In that case, the average power expended could be readily computed from the work computed in Equation (S14) by taking the temporal average of the integrand, *i.e.*,

$$P(\mathbf{x},t) = \frac{W(\mathbf{x},t)}{t}. \tag{S16}$$

During epithelial expansion, however, the edges of the tissue move to invade new space, meaning that some spatial locations will only contain tissue for a subset of the experimental time. This implies that dividing through by the total time, as in Equation (S16), will bias the average power expended by the tissue at the tissue edges. We address this issue by correcting for the total time that any given spatial location has been occupied by epithelium to write

$$P(\mathbf{x},t) = \frac{\int_0^t \mathbb{I}_{\text{tissue}}(\mathbf{x},s) W(\mathbf{x},s) \mathrm{d}s}{\int_0^t \mathbb{I}_{\text{tissue}}(\mathbf{x},s) \mathrm{d}s}. \tag{S17}$$

Here, $\mathbb{I}_{\text{tissue}}(\mathbf{x},s)$ is an indicator function that is equal to one if there is tissue present at location $\mathbf{x}$ at time $s$ and zero otherwise. Note that Equation (S17) reduces to Equation (S16) when $\mathbb{I}_{\text{tissue}}(\mathbf{x},s) \equiv 1$, which is particularly true in the bulk, *i.e.*, the centre of the tissue, which is always populated. The general formulation in Equation (S17) provides an unbiased quantity that compares the time-averaged rate of mechanical work at any spatial location in the tissue and at any time. Finally, we remark that this average power metric takes into account the average rate of work from the beginning of the experiment, and so all of past time contributes equally to this metric.

### COMPUTATION OF STREAMLINES AND LOCAL VELOCITIES

For the computation of each regulation space, we consider a dataset that consists of points within a given spatiotemporal domain, *e.g.* all spatial locations between $t = 1$h and $t = 3$h for a 3V/cm pulse. We then computed the time derivative of the quantities of velocity and power using a first-order finite difference scheme.

The time derivatives of velocities and power can be used directly to compute velocities within $v$-$p$ space, as the regulation space is a vector field, $R$, where

$$R(x,t) = \frac{\mathrm{d}}{\mathrm{d}t} \left( v(x,t), p(x,t) \right). \tag{S18}$$

We graphically produced regulation spaces using the MATLAB inbuilt function streamslice.

### COMPUTATION OF CRITICAL POWER

Having computed power for each spatial location at each given time point, across the different biological replicates, we make a mesh grid of 50 points interpolating between the bottom 5% and top 95% recorded power in the dataset. For each point, we compute the mean and standard deviation of the computed jerk at that time and spatial location. Plotting these quantities against each other gives a scatter plot of binned power against binned jerk. To determine the critical power, we then fit a piecewise linear spline between the mean datapoints and determine $p_0$ as the first point of intersection with the zero-jerk line.

### COMBINING SINGLE CELL TRACKING AND PIV

#### Backtracking

Since the mechanical model in Equation (S17) leads to a continuum metric of average power, we need to relate the power as computed in Equation (S17) to individual cells, depending on their position in time and space. This is done as follows. For each cell track, we use the location of the cell nucleus to compute the value of the mechanical power at that location, using linear interpolation of the power computed using Equation (S17). Recall that the time-averaged power was computed using the material derivative, so each cell is assigned the average mechanical power of the tissue parcel it is contained in. For transparency, we note that the dimension of the PIV window corresponds to roughly three cell lengths. With each track now consisting of a location, velocity, and average mechanical power, we can explore the joint behaviours of velocity and power in single cells.

#### Switching time analysis

To assess how long single cells remain in a state of being high or low spenders, we assess the probability density function of the time needed to make a switch. To explore this, cells were classified as *high spenders* if their time-averaged mechanical power was higher than the average in the monolayer (and *low spenders* if lower). For fair comparisons between experimental conditions, all analysis was done for observations between $t = 1$ hour and $t = 3$ hours, *i.e.*, the times that correspond to the EF being turned on during the electrotaxis experiment. Additionally, in order to obtain an unbiased estimate of the switching probabilities between being in a high spending state or in a low spending state, we restrict our analysis to the tracks that contain cell locations for the entire stimulation window. As a first step to understand the dynamics of switching from a high to a low spending state, we computed the time that each cell spent between two switching events. These *residence times* show a striking empirical density, as shown in Figure **Fig. S1**.

The empirical distributions display fat tails, which suggests that the probability density of the residence times might be more similar to a power law than to an exponential distribution. To investigate the distribution of the residence times, the histograms of the residence times can be compared to a least squares fit of an exponential distribution, whose probability density function (pdf) is given by

$$f(x) = \lambda e^{-\lambda x}, \quad x > 0, \tag{S19}$$

where $\lambda$ is the *rate*. We also consider a power law distribution, whose distribution is given by

$$f(x) = a x^{a-1}, \quad x > 0, a < 1, \tag{S20}$$

where $a$ is the *exponent*. It can be seen in Figure **Fig. S1** that the power law distributions are in good agreement with the empirical distributions of the residence times, whereas the exponential distributions underestimate the probabilities that cells remain in the same state for a long times. To assess the goodness of fit of the power law distribution to the data, and to understand whether there is a good agreement with the data, we plot the empirical densities obtained from the histograms in Figure **Fig. S1** on double-log axes in Figure **Fig. S2**. The log-log plots in Figure **Fig. S2** demonstrate that there is excellent agreement between the empirical probability density of the residence times and the fitted power law distributions. Moreover, we find that the exponents for the least-squares fit to the empirical distributions of the residence times arising from the different stimulation traces and the control experiment are all very similar (Table **Table S2**). Strikingly, all distributions, regardless of EF strength, follow a very similar distribution, suggesting that the temporal dynamics of switching spending state are agnostic to external stimuli. This is surprising because it suggests that the EF stimulation protocol changes how much time-averaged power cells change, but it does not change how cells collectively organise their power

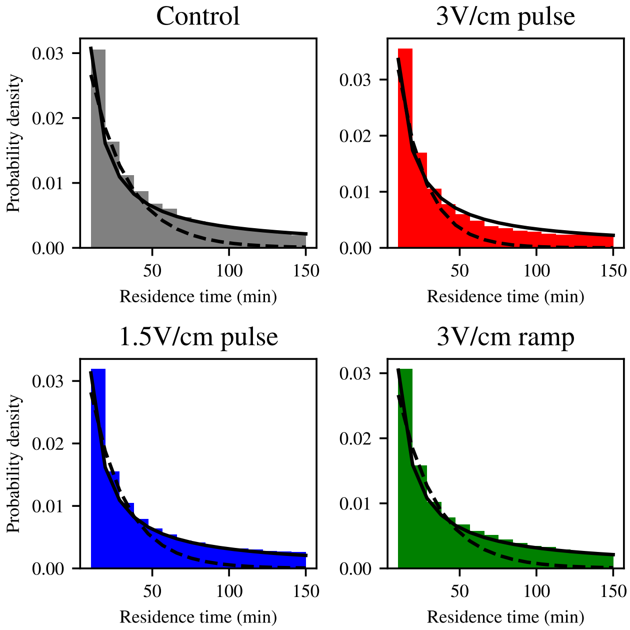


**Fig. S1.** Empirical probability density function of the lengths of times spent being either a high or a low spender for each of the experiments, compared to linear squares fit for exponential distribution (dashed line) or power law distribution (solid line).

| Experiment | Computed exponent |
|---|---|
| No stim (control) | 0.325 |
| 3 V/cm pulse | 0.336 |
| 1.5 V/cm | 0.313 |
| 3 V/cm ramp | 0.319 |

**Table S2.** Computed power law exponents for the switching times pdf in the different experiments.

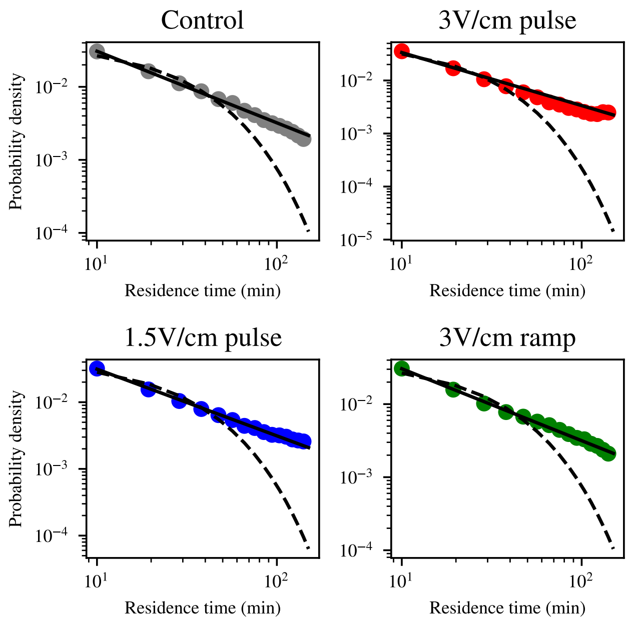


**Fig. S2**. Log-log plot of empirical probability densities for the lengths of times spent being either a high or a low spender (scatter), compared to linear squares fit for exponential distribution (dashed line) or power law distribution (solid line).

expenditure. Since therefore the mechanism underlying this collective regulation appears unchanged between different EF stimulation protocols, the similarity between the distributions of the residence times points to mechanical factors inherent to epithelial migration, such as cell shape index and tissue fluidity, playing a role in influencing state switching. Finally, the fact the power law is a good fit to the empirical distribution of the times spent in either of the energetic states, and the exponential law is not, suggests that state switching is a random process with some memory in the system, as has previously been suggested in electrotaxing epithelia [15].

‡
§

[1] D. T. Tambe, U. Croutelle, X. Trepat, C. Y. Park, J. H. Kim, E. Millet, J. P. Butler, and J. J. Fredberg, PLOS ONE **8**, 1 (2013), URL https://doi.org/10.1371/journal.pone.0055172.
[2] R. Alert, C. Blanch-Mercader, and J. Casademunt, Phys. Rev. Lett. **122**, 088104 (2019), URL https://link.aps.org/doi/10.1103/PhysRevLett.122.088104.
[3] R. Alert and X. Trepat, The Annual Review of Condensed Matter Physics is Annu. Rev. Condens. Matter Phys. 2020 **11**, 77 (2020), URL https://doi.org/10.1146/annurev-conmatphys-.
[4] X. Serra-Picamal, V. Conte, R. Vincent, E. Anon, D. T. Tambe, E. Bazellieres, J. P. Butler, J. J. Fredberg, and X. Trepat, Nature Physics **8**, 628 (2012), ISSN 17452481.
[5] C. Blanch-Mercader, R. Vincent, E. Bazellières, X. Serra-Picamal, X. Trepat, and J. Casademunt, Soft Matter **13**, 1235 (2017), URL http://dx.doi.org/10.1039/C6SM02188C.
[6] K. Vazquez, A. Saraswathibhatla, and J. Notbohm, Scientific Reports **12** (2022), URL https://doi.org/10.1038/s41598-022-06504-0.
[7] S. Tlili, E. Gauquelin, B. Li, O. Cardoso, B. Ladoux, H. D. Ayari, and F. Graner, Royal Society Open Science **5** (2018), ISSN 20545703.
[8] S. Tlili, M. Durande, C. Gay, B. Ladoux, F. Graner, and H. Delanoë-Ayari, Physical Review Letters **125** (2020), ISSN 10797114.
[9] A. Christensen, A.-K. V. West, L. Wullkopf, J. T. Erler, L. B. Oddershede, and J. Mathiesen, Physical Biology **15**, 066004 (2018), URL https://dx.doi.org/10.1088/1478-3975/aacedc.
[10] C. Fang, J. Yao, Y. Zhang, and Y. Lin, Biophysical Journal **121**, 1266 (2022), ISSN 0006-3495, URL https://www.sciencedirect.com/science/article/pii/S0006349522001527.
[11] J. C. Arciero, Q. Mi, M. F. Branca, D. J. Hackam, and D. Swigon, Biophysical Journal **100**, 535 (2011), ISSN 15420086.
[12] C. Pérez-González, R. Alert, C. Blanch-Mercader, M. Gómez-González, T. Kolodziej, E. Bazellieres, J. Casademunt, and X. Trepat, Nat. Phys. **15**, 79 (2019).
[13] S.-Z. Lin, P.-C. Chen, L.-Y. Guan, Y. Shao, Y.-K. Hao, Q. Li, B. Li, D. A. Weitz, and X.-Q. Feng, Advanced Biosystems **4**, 2000065 (2020), https://onlinelibrary.wiley.com/doi/pdf/10.1002/adbi.202000065, URL https://onlinelibrary.wiley.com/doi/abs/10.1002/adbi.202000065.
[14] S.-Z. Lin, W.-Y. Zhang, D. Bi, B. Li, and X.-Q. Feng, Communications Physics **4**, 21 (2021), ISSN 2399-3650, URL https://doi.org/10.1038/s42005-021-00530-6.
[15] A. E. Wolf, M. A. Heinrich, I. B. Breinyn, T. J. Zajdel, and D. J. Cohen, PNAS Nexus **1** (2022).
