## Supplemental Figures for "Cellular cruise control: energy expenditure as a regulator of collective migration in epithelia"

**Supplementary Movies**

**Supplementary Video 1:** 10x phase contrast video of a freely expanding 5x5mm MDCK-II epithelium. 10 min/frame. Scale bar = 500um.

**Supplementary Video 2:** Average Speed [um/h] heatmap of freely expanding 5x5mm MDCK-II epithelia, averaged over all replicates. 10 min/frame.

**Supplementary Video 3:** Average Traction [Pa/um] heatmap of freely expanding 5x5mm MDCK-II epithelia, averaged over all replicates. 10 min/frame.

**Supplementary Video 4:** Average Work [J/m] heatmap of freely expanding 5x5mm MDCK-II epithelia, averaged over all replicates. 10 min/frame.

**Supplementary Video 5:** Average Power [W/m^2] heatmap of freely expanding 5x5mm MDCK-II epithelia, averaged over all replicates. 10 min/frame.

**Supplementary Video 6:** 10x phase contrast video of a 5x5mm MDCK-II epithelium undergoing electrotaxis. 1hr 0 V/cm, 3hr 3 V/cm, 4hr 0 V/cm. 10 min/frame. Scale bar = 1000um.

**Supplementary Video 7:** 10x phase contrast video of a portion of an electrotaxing epithelium, magnified for visualization purposes. 1hr 0 V/cm, 3hr 3 V/cm, 4hr 0 V/cm. 10 min/frame. Scale bar = 250um.

**Supplementary Video 8:** Average Power [W/m^2] heatmap of electrotaxing 5x5mm MDCK-II epithelia, averaged over all replicates. 1hr 0 V/cm, 3hr 3 V/cm, 4hr 0 V/cm. 10 min/frame.

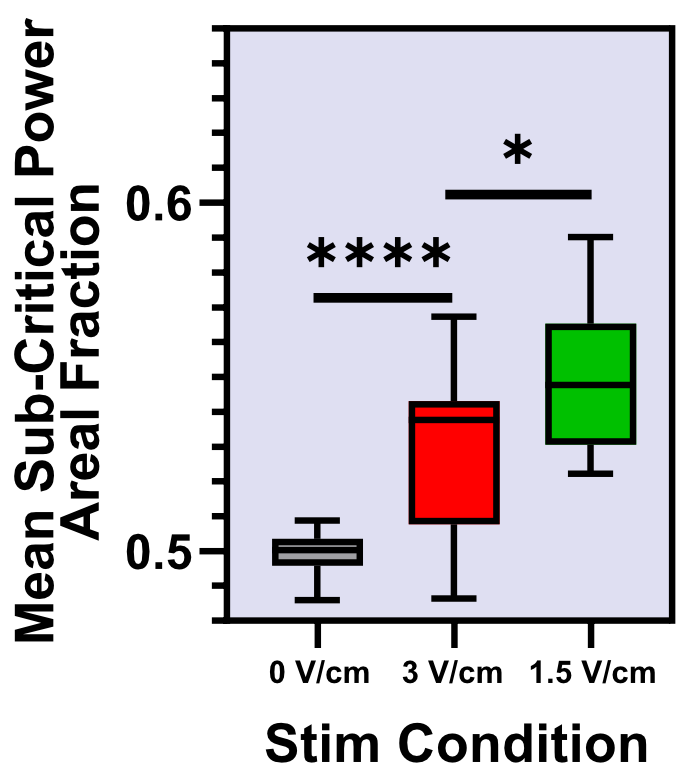

**Supplementary Figure 1:** Average areal fraction of positive jerk *during* stimulation, averaged over all tissues for different stimulation traces.

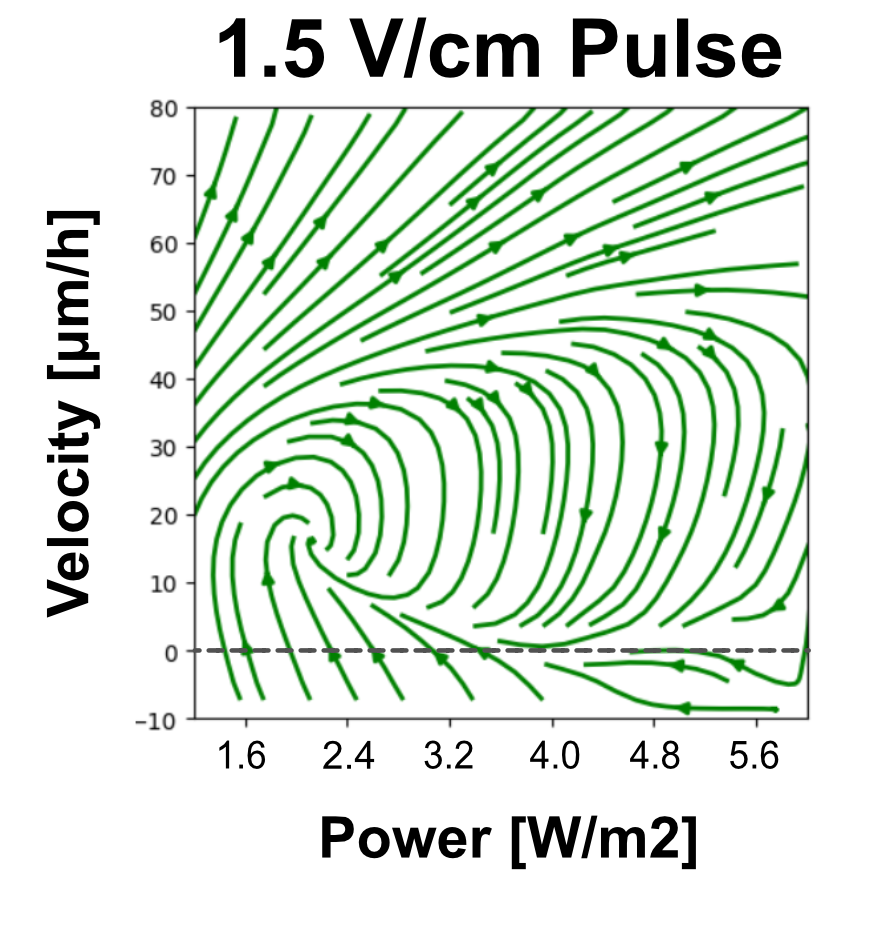

**Supplementary Figure 3:** v-p space averaged over all tissues stimulated with a 1.5 V/cm pulse.

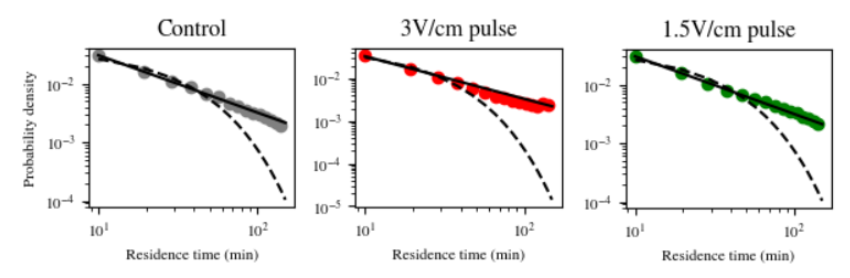

**Supplementary Figure 3:** Probability density distributions of residence times of cells in a high or low spending regime. Points represent averages over all cells. Straight line represents a power law, dotted line represents an exponential fit.

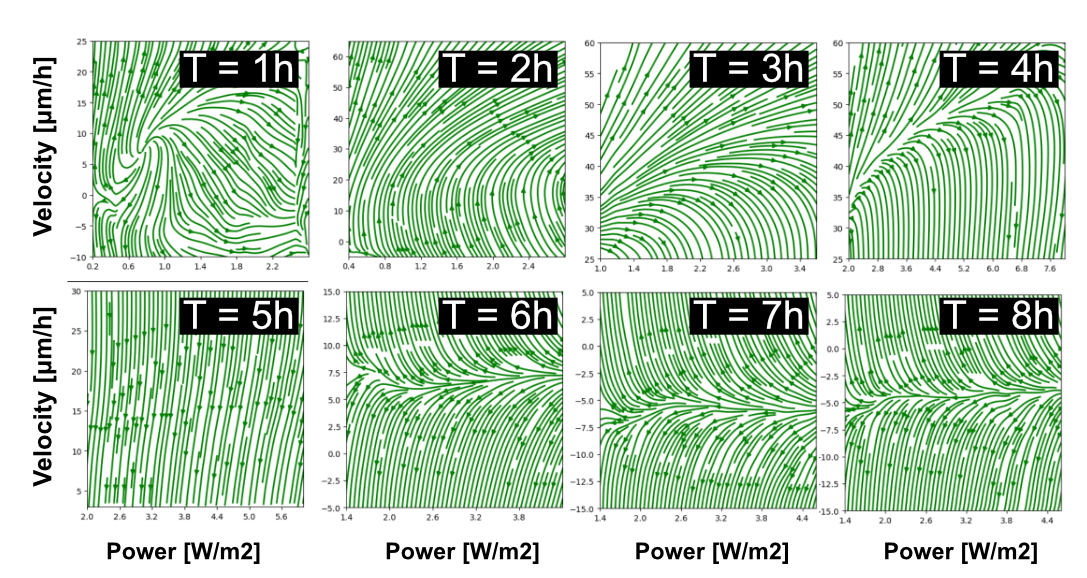

**Supplementary Figure 4:** Phase spaces for one-hour intervals averaged over all 1.5 V/cm stimulated tissues. Note the important changes in axis range between timepoints.
